# Potential involvement of root auxins in drought tolerance by modulating nocturnal and daytime water use in wheat

**DOI:** 10.1101/530246

**Authors:** Walid Sadok, Rémy Schoppach

**Affiliations:** Department of Agronomy and Plant Genetics, 1991 Upper Buford Circle, 411 Borlaug Hall, University of Minnesota, St. Paul, MN 55108-6026, USA; Earth and Life Institute, Université Catholique de Louvain, Louvain-la-Neuve 1348, Belgium

**Keywords:** Drought tolerance, water-saving, transpiration rate, nocturnal, nighttime, auxin, abscisic acid, root hydraulics, vasculature, wheat, yield

## Abstract

The ability of wheat genotypes to save water by reducing their transpiration rate (TR) under times of the day with high vapour pressure deficit (VPD) has been linked to increasing yields in terminal drought environments. Further, recent evidence shows that reducing nocturnal transpiration (TR_N_) could amplify water-saving. Previous research indicates that such traits involve a root-based hydraulic limitation, but the contribution of hormones, particularly auxin and abscisic acid (ABA) has not been explored to explain the shoot-root link. In this investigation, based on physiological, genetic and molecular evidence gathered on a mapping population, we hypothesized that root auxin accumulation regulates whole-plant water use during both times of the day. Eight double-haploid lines were selected from a mapping population descending from two parents with contrasted water-saving strategies and root hydraulic properties. These spanned the entire range of slopes of TR responses to VPD and TR_N_ encountered in the population. On those lines, we examined daytime/night-time auxin and ABA contents in the roots and the leaves in relation to hydraulic traits that included whole-plant TR, plant hydraulic conductance (K_Plant_), slopes of TR responses to VPD and leaf-level anatomical traits. Root auxin levels were consistently genotype-dependent in this group irrespective of experiments and times of the day. Daytime root auxin concentrations were found to be strongly and negatively correlated with daytime TR, K_Plant_ and the slope of TR response to VPD. Night-time root auxin levels significantly and negatively correlated with TR_N_. In addition, daytime and night-time leaf auxin and ABA concentrations did not correlate with any of the examined traits. The above results indicate that accumulation of auxin in the root system reduces daytime and night-time water use and modulates plant hydraulic properties to enable the expression of water-saving traits that have been associated with enhanced yields under drought.

## INTRODUCTION

Mediterranean-type terminal drought events are the most penalizing water deficit regimes experienced by crops and their frequency and intensity is expected to increase across the globe as a result of anthropogenic climate change (Berger *et al*., 2016). Under such conditions, plants typically grow on stored soil moisture, following early-season precipitation which typically triggers planting from the farmer. In those environments, plants will have to achieve a seed-to-seed cycle by relying on an amount of stored soil water that did not evaporate, runoff or percolate to deeper, inaccessible soil layers. For subsistence farming, the situation is even more challenging since the crop will have to generate yields that have to be economically viable for the household (Solh and Van Ginkel, 2014).

Water-saving traits have a promising potential for enhancing yields under terminal drought environment (e.g., Sinclair *et al*., 2017). Over the last decade, a series of simulation studies using physiologically-informed, process-based simulation modelling consistently demonstrated for several crops such as maize (Zea *mays* (L.)), soybean (*Glycine max* (L.) Merr.) and lentil (*Lens culinaris* (Medik.)) that the expression of such traits would translate into in increased probability of significant yields benefits (e.g., Sinclair *et al*., 2010; Messina *et al*., 2015; Guiguitant *et al*., 2017). In those studies, a functional trait that was found to consistently generate yield benefits under terminal drought consisted in decreased transpiration rates (TR) during times of the day where the levels of atmospheric vapour pressure deficit (VPD) would outmatch the ability of the plant to supply water to the transpiring leaf, resulting in a decrease in canopy conductance (Sinclair *et al*., 2005; Sinclair *et al*., 2017).

On wheat (*Triticum aestivum* (L.)), such trait has been shown to be consistently expressed among historical genotypes released for the rain-fed production environments in Australia between 1890 and 2008 (Schoppach *et al*., 2017). Experimentation carried out on an elite, drought-tolerant Australian breeding line ‘RAC875’ showed that this water-saving behaviour was stemming from a lower root hydraulic conductivity that was not exhibited by a drought-sensitive check cultivar called Kukri (Schoppach *et al*. 2012; Schoppach *et al*. 2014a). This difference was traced to an increased resistance to radial, trans-membrane water movement in the roots of RAC875, putatively controlled by a lack of a mercury-sensitive aquaporin population (Schoppach *et al*., 2014a). In addition, the roots of RAC875 were found to exhibit particularly smaller diameters of central metaxylem (CMX) vessels, below a limit (55 μm) that was found by Richards and Passioura (1989) to be associated the with up to 11% of yield increases in a breeding program that specifically targeted decreasing CMX diameters to achieve water-saving in the field in Australia.

Currently, there is evidence supporting the idea that whole-plant TR response to VPD involves an interaction between ‘local’ (i.e., leaf-based) and ‘non-local’ (root-based) mechanisms mobilizing long-distance hydraulic signals that involve a combination of root anatomical traits and a dynamic control over radial water flow mediated by aquaporins (Vandeleur *et al*., 2014; Vadez, 2014; Maurel *et al*., 2016; Sivasakthi *et al*., 2017). However, other investigations point to a role played by a shoot-to-root hormonal signal, namely abscisic acid (ABA) in coordinating TR with root hydraulic conductivity in response to increasing VPD (Kudoyarova *et al*., 2011; Veselov *et al*., 2018).

In an effort to illuminate clues underpinning these links, a high-throughput phenotyping approach for characterizing TR response curves to naturally increasing VPD was undertaken in a double-haploid (DH) mapping population resulting from a cross between RAC875 and Kukri (Schoppach *et al*., 2016). The genetic analysis revealed a major QTL controlling these responses, which explained more than 25% of the genetic variance, with a peak region that was mapped to 9 drought-tolerance candidate genes that were found to be expressed in the roots independent from phenology, in a RNA-Seq experiment (Schoppach *et al*., 2016). In further support of the root-based origin of these responses, the putative functions of several of those genes directly indicated their involvement in root development, root xylem patterning and response to ABA and water stress. However, an unexpected finding of that study was that several of the genes were also auxin-related, raising the speculation that root auxin could be directly involved in the variation in TR responses to VPD found in this population.

Considering evidence documenting direct involvement of auxin i) in root development and vasculature patterning (Fàbregas *et al*., 2015; Alabdallah *et al*., 2017), ii) in down-regulating aquaporin (AQP) expression and hydraulic conductivity (Péret *et al*., 2012) and iii) as a shoot-root signalling molecule (Vandeleur *et al*., 2014), a first goal of this research was to examine the involvement of root auxin levels in variation of traits associated with TR responses to VPD. To this end, we examined the relationship between root auxin concentrations and variation in whole-plant hydraulic conductance, daytime TR and TR response curves to increasing VPD among a group of 8 DH lines from the RAC875 x Kukri cross, which previously expressed variation in canopy conductance spanning the entire range of the population.

A second goal of the investigation was to evaluate two alternative, competing hypotheses to the direct involvement of root auxin in regulating whole-plant hydraulics in wheat. The first alternative hypothesis is based on the idea that leaf auxin could be involved in regulating those hydraulic traits. Considering findings documenting the role played by leaf auxins in leaf vasculature development, particularly xylem (Taneda and Terashima, 2012; Moreno-Piovano *et al*., 2017) and given that leaf vasculature plays a central role in leaf hydraulic conductance (Caringella *et al*., 2015), we examined the relationship between leaf auxin concentrations, the above hydraulic traits, and a group of six vasculature-related, leaf anatomical traits. The second alternative hypothesis consisted of the involvement of shoot and/or root ABA in regulating those traits. Such possibility is supported by findings of Kudoyarova *et al*. (2011) on durum wheat and Veselov *et al*. (2018) on barley, indicating that high VPD triggers leaf ABA export to the root to regulate root hydraulic conductivity, as a way to maintaining leaf hydration as increases. In contrast, the findings of Kholovà *et al*. (2010) on pearl millet indicate a localized phenomenon where the accumulation of leaf ABA under high VPD drives a decrease in stomata conductance and therefore the expression of water-saving of drought-tolerant genotypes. To address these hypotheses, another goal of the investigation was to examine the relationships between root and leaf ABA and whole-plant hydraulic conductance, daytime TR and TR response curves to increasing VPD among the same eight lines selected from the RAC875 x Kukri population.

In addition to traits controlling daytime water use, there is indirect evidence to suggest potential yield benefits arising from reduced nocturnal transpiration in crops grown in drought-prone environments. In the case of Australian wheat, Rawson and Clarke (1988) found that night-time transpiration rates (TR_N_) could be in excess of 0.5 mm per night, first hypothesizing potential yield benefits resulting from night-time water-saving. This potential was confirmed in a series of studies on crops including wheat (Schoppach *et al*., 2014b) bean (Resco de Dios *et al*., 2015) and grapevine (Coupel-Ledru *et al*., 2016). Under controlled environment conditions, Schoppach *et al*., (2014b) identified significant genotypic variability in wheat TR_N_, which was driven by nocturnal VPD, reaching values that were up to 55% of maximal daytime TR, under high levels of nocturnal VPD (2.1 kPa). Interestingly, those responses were found to mirror daytime TR response curves to VPD, in that the water-saving genotype RAC875 also exhibited a particularly limited TR_N_ under high nocturnal VPD, in contrast to the drought-sensitive cultivar Kukri. More recently, Claverie *et al*. (2018) found that under water-deficit conditions, TR_N_ in wheat was much less sensitive to progressive soil drying relative to daytime TR, resulting in a progressively higher contribution of TR_N_ to daily water use. In that study, RAC875 was found to exhibit a tighter control of TR_N_ than Kukri during the soil drying sequence, resulting in the expression of what could result in a water-saving behaviour (Claverie *et al*., 2018).

While the genetic basis of TR_N_ was found to be controlled by numerous QTL in the RAC875 x Kukri population (Schoppach *et al*., 2016), the physiological basis driving this variation remains poorly understood. On well-watered wheat, Claverie *et al*. (2016) found that high TR_N_ levels were associated with changes in root anatomy, particularly smaller endodermis cell size and reduced xylem sap exudation rates, a proxy measure for root pressure. On trees, high TR_N_ levels have been associated with higher N uptake by the roots to compensate for low N availability (Rohula *et al*., 2014), or with the need for higher O_2_ delivery via the nocturnal xylem sap to maintain functions sustained via dark respiration (Marks and Lechowicz, 2007). However, so far, hormonal involvement in regulating TR_N_ was not explored. Given the involvement of root auxin in regulating root vasculature development (Fàbregas *et al*., 2015; Alabdallah *et al*., 2017), and nitrogen foraging by the roots (Song *et al*., 2013), we examined in this investigation the relationship between root auxin concentration and TR_N_ levels among the same 8 DH lines from the RAC875 x Kukri cross. Similar to the first goal of this investigation, we also tested the hypothesis of the involvement of night-time leaf auxin levels and leaf anatomical traits in controlling TR_N_. Finally, we examined the possibility that night-time leaf or root ABA could be involved in the variation in TR_N_ in this group.

## MATERIALS AND METHODS

### Genetic material

Eight genotypes were selected from a population of 143 bread wheat (*Tricticum aestevium* (L.)) double haploid lines that descended from a cross between the drought-tolerant breeding line RAC875 (RAC655/3/Sr21/4*LANCE//4*BAYONET) and the check, drought-sensitive cultivar Kukri (76ECN44/76ECN36//MADDEN/6*RAC177). The 8 genotypes were selected such that they span the entire range of the slopes of TR response curves to increasing VPD that was found for this population (Schoppach *et al*., 2016; [Supplementary Data Fig. S1]).

### Growth conditions

Four independent experiments were undertaken in this study (Table 1). In all experiments, plants were grown for 34-37 days in a glasshouse at the Université catholique de Louvain, Belgium (50°40’N, 4°36’E). The glasshouse was equipped with a supplementary LED lighting system (Pro650, LumiGrow, Novato, California, USA), which provided an additional photosynthetic photon flux density (PPFD) of 200 μmol m^−2^ s^−1^ to the natural ambient PPFD at canopy level. The lighting system activated each time the incident radiation dropped below 500 W m^−2^ between 0600 h am and 2200 h in experiments E1 and E2 and 0800 h and 2000 h in experiment E3. Plants were sown and grown in well-watered garden soil (DCM Corporation, Grobendonk, Belgium) in experiment E1 and in a hydroponic system in experiments E2 and E3 (see below for details).

**Table 1.** Summary of the experiments.

| Experiment | Sowing date | Measurement Date | Measured variable | Plant age (days) | Root medium | Growth period conditions (± S.E.) | | | |
| --- | --- | --- | --- | --- | --- | --- | --- | --- | --- |
|  |  |  |  |  |  | Daytime |  | Nighttime |  |
| | | | | | | T ($^{\circ}$C) | VPD (kPa) | T ($^{\circ}$C) | VPD (kPa) |
| E1 | 28/04/15 | 02/06/15 | Hydraulic conductance | 36 | Potting mix | $27.6 \pm 0.4$ | $2.6 \pm 0.10$ | $21.1 \pm 0.2$ | $1.4 \pm 0.03$ |
| E2 | 12/05/15 | 17/06/15 | Nighttime IAA and ABA, TR<sub>N</sub> | 37 | Hydroponic | $28.4 \pm 0.5$ | $2.8 \pm 0.12$ | $21.5 \pm 0.3$ | $1.4 \pm 0.03$ |
| E3.1 | 21/12/16 | 24/01/17 | Nighttime IAA and ABA, TR<sub>N</sub> | 34 | Hydroponic | $25.8 \pm 0.1$ | $1.6 \pm 0.03$ | $24.6 \pm 0.1$ | $1.7 \pm 0.02$ |
| E3.2 | 21/12/16 | 24/01/17 | Nighttime IAA and ABA, TR<sub>N</sub> | 34 | Hydroponic | $25.8 \pm 0.1$ | $1.6 \pm 0.03$ | $24.6 \pm 0.1$ | $1.7 \pm 0.02$ |
| E3.3 | 21/12/16 | 24/01/17 | Daytime IAA and ABA, TR | 34 | Hydroponic | $25.8 \pm 0.1$ | $1.6 \pm 0.03$ | $24.6 \pm 0.1$ | $1.7 \pm 0.02$ |
| E4 | 07/05/14 | 12/06/14 | Leaf anatomical traits | 36 | Potting mix | $28.6 \pm 0.5$ | $2.2 \pm 0.10$ | $22.8 \pm 0.3$ | $1.3 \pm 0.03$ |

Potted plants were grown as described in Schoppach *et al*. (2016). Briefly, three seeds per pot were sown at a depth of 2.5 cm in custom-made PVC columns (0.11 m diameter and 0.33 m tall) filed with 1205 ± 5g of garden soil. Ten days after sowing, each pot was thinned to a single plant. Pots were watered regularly until the measurements (hydraulic conductance, see below) were initiated. Hydroponically-grown plants (E2 and E3) were grown as in Schoppach *et al*. (2014a) with the exception that plant growth took place in the same greenhouse and under relatively similar conditions as the potted plants (Table 1). Seeds were germinated for 3 days in petri dishes on a layer of filter paper humidified with ultra-pure water, inside a dark climate cabinet where temperature (T) was maintained at 20 °C. The germinated seeds were subsequently placed on expanded polystyrene plates (30 plants per plate) floating on a nutrient solution that filled 26-L plastic tanks. The solution was regularly aerated by means of an automatic pump system (flow rate: 150 L/h, for 15 min every hour) and was prepared based on the method described by Rengel and Graham (1996). This solution contained: 2mM Ca(NO3)2, 0.5mM MgSO4.7H2O, 1.5mM KNO3, 0.1mM KC1, 2mM MES-KOH, 0.1mM NH4H2PO4, 10mM H3BO3, 0.1 mM Na2 MoO4, 25mM K3-(N-(2-hydroxyethyl) ethylenedinitrilotriacetic acid) (HEDTA), 0.1mM FeHEDTA, 1 mMMnHEDTA, 0.5 mMCuHEDTA, 0.1 mMNiHEDTA, 2 mM ZnHEDTA. During the first 6 days, seedlings were grown in a half-strength nutrient solution. The solutions were replaced every week. The pH of the nutrient solution was checked daily and when necessary adjusted to a value of 6 using MES-KOH. During all experiments, T and relative humidity (RH) conditions were continuously recorded every five minutes by pocket sensors connected to USB dataloggers (EL-USB-2-LCD, Lascar Electronics, Whiteparish, United Kingdom) placed in 3-5 locations across the setup.

### Plant hydraulic conductance

This experiment (E1) was carried out on the potted plants (Table 1). On the day prior to the measurements, three replicate plants per genotype (i.e., a total of 24 pots) were watered to dripping at 1600 h. They were slightly re-watered again on the following morning around 0500 h to minimize the occurrence of transient soil moisture deficit during the measurements. Afterwards, the soil in each pot was covered with an aluminium foil to nullify direct soil water evaporation. The plants were then placed inside a walk-in growth chamber (PGV36, Conviron, Winnipeg, Manitoba, Canada) and left to acclimate for 6 hours under the steady-state conditions Photosynthetic Photon Flux Density (PPFD) = 480 μmol m^−2^ s^−1^, T = 31.9 °C, VPD = 2.8 kPa, Table 2).

**Table 2.**
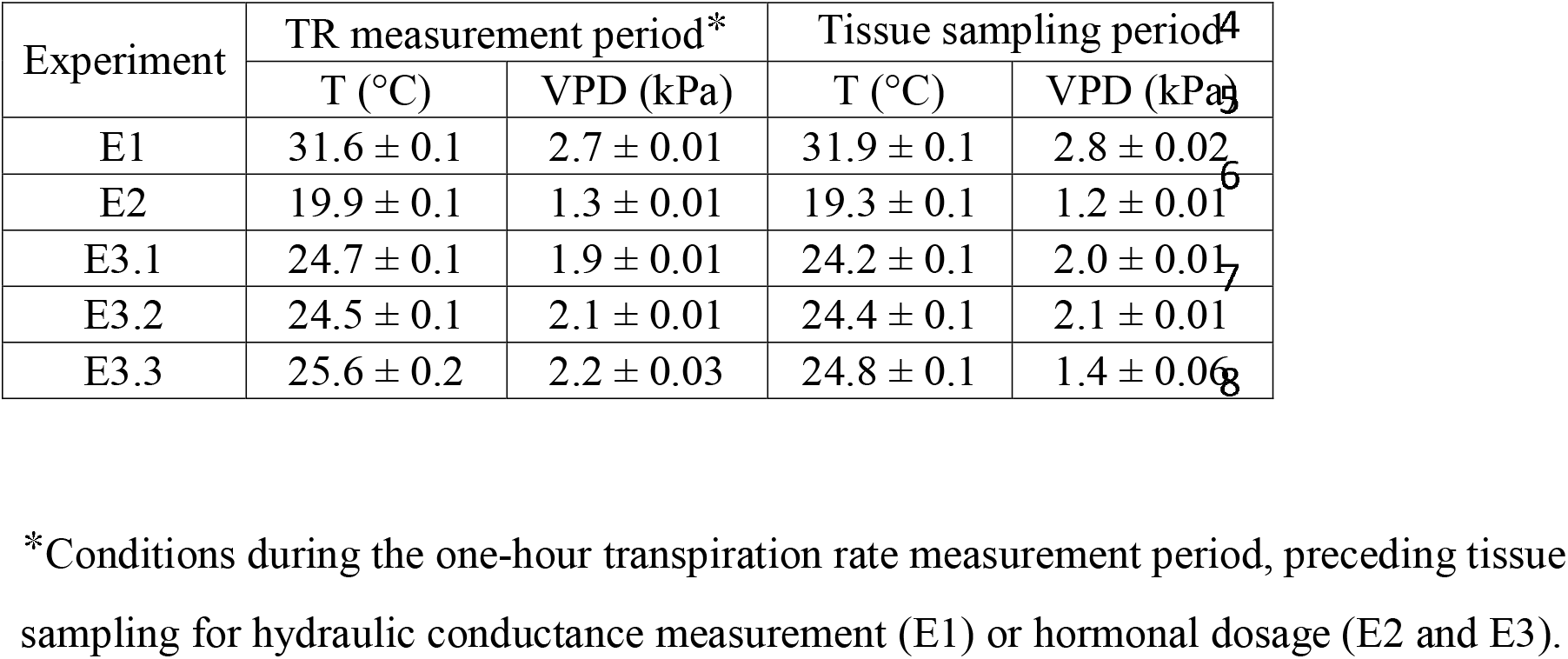
Environmental conditions during the measurements.

| Experiment | TR measurement period* |  | Tissue sampling period<sup>4</sup> |  |
| --- | --- | --- | --- | --- |
|  | T (°C) | VPD (kPa) | T (°C) | VPD (kPa) |
| E1 | 31.6 ± 0.1 | 2.7 ± 0.01 | 31.9 ± 0.1 | 2.8 ± 0.02 |
| E2 | 19.9 ± 0.1 | 1.3 ± 0.01 | 19.3 ± 0.1 | 1.2 ± 0.01 |
| E3.1 | 24.7 ± 0.1 | 1.9 ± 0.01 | 24.2 ± 0.1 | 2.0 ± 0.01 |
| E3.2 | 24.5 ± 0.1 | 2.1 ± 0.01 | 24.4 ± 0.1 | 2.1 ± 0.01 |
| E3.3 | 25.6 ± 0.2 | 2.2 ± 0.03 | 24.8 ± 0.1 | 1.4 ± 0.06 |

\*Conditions during the one-hour transpiration rate measurement period, preceding tissue sampling for hydraulic conductance measurement (E1) or hormonal dosage (E2 and E3).

Following the acclimation period, whole-plant TR values were determined for all 8 lines by performing 2 weightings separated by 60 min, using an electronic balance with a resolution of 0.01 g (Model Fx-3000i, A&D Co. Ltd, Tokyo, Japan). TR (mg H_2_O m^−2^ s^−1^) was later calculated as the difference in pot mass, normalized by whole plant leaf area which was measured destructively, using a leaf area meter (LI-3100C, Li-Cor, Lincoln, NE, USA). Leaf water potential (□_leaf_) of the uppermost fully developed leaf was measured using a standard pressure chamber (Scholander 670, OMS Instrument, Albany, USA). Whole plant conductance (K_Plant_) was defined as the flux of water through the plant (TR), divided by the water potential gradient between the soil (□_soil_) and the top leaf (□_leaf_), as follows:

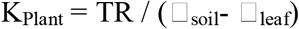

Since the pots were well-watered, it was assumed that □_soil_ is approaching 0 MPa. Consequently, the unit of K is: mg H_2_O leaf area m^−2^ s^−1^ MPa^−1^.

### Auxin and abscisic acid measurements

#### Tissue harvest conditions

Measurements were carried out on hydroponically-grown plants (E2 and E3, Table 1). Prior to measurements, all plants measured in E2 and E3 were cautiously removed from the polystyrene plates at the end of the afternoon and transferred into individual 300-mL dark brown glass bottles covered with aluminium foil and filled with a fresh hydroponic solution at 2000 h. Plants were then placed in the glasshouse at a PPFD of 0 μmol m^−2^ s^−1^ under the environmental conditions displayed in Table 2.

Nocturnal auxin (indole-3-acetic acid or IAA) and ABA measurements were made on leaf and root tissues (4 replicate plants per genotype) sampled at 0400 h in experiment E2 and at two times of the night (E3.1: 0200 h and E3.2: 0500 h) in experiment E3, under conditions where PPFD was zero (Table 2). Daytime measurements of leaf and root IAA and ABA (experiment E3.3) were made on tissues sampled in the morning at 0800 h, under conditions reported in Table 2. Prior to each one of these samplings, whole-plant TR was determined gravimetrically during the previous hour using the same approach as mentioned earlier. In total, this resulted in 12 and 4 nocturnal TR (TR_N_) and daytime TR measurements per genotype, respectively. Temperature and VPD conditions observed during this period are reported on Table 2.

For leaf hormonal dosage, a predetermined segment located in the middle of the uppermost fully expanded leaf was quickly cut using a sharp scalpel blade and flash-frozen in liquid nitrogen. The leaf area represented by the cut leaf segment was accounted for when normalizing calculating whole plant transpirational water loss by leaf area. Immediately after leaf harvesting, whole root systems were harvested for hormonal dosage by de-rooting the plant using a sharp blade, and mopping the roots quickly on water-absorbing tissue before placing them inside falcon tubes which were flash-frozen in liquid nitrogen.

#### Hormone extraction and dosage

After harvesting, leaf and root samples were ground in liquid nitrogen, homogenized, packaged in Eppendorf tubes and stored in a −80°C freezer. Free IAA and ABA were extracted based on the extraction procedure described in Prinsen *et al*., (2000). Briefly, homogenized plant material (60-80 mg) was extracted overnight in 80% methanol (10 μl/mg fresh weight). C613-phenyl-IAA (50 pmol, Cambridge Isotope Laboratories Inc., Andover, Massachusetts, USA) and D6-ABA (100 pmol, (±)-3’, 5′, 5′, 7′, 7′, 7′-d6 ABA, National research council Canada, Saskatoon, Canada) were added as internal standards. After centrifugation (20000 g, 15 min, 4 °C, 5810R, rotor FA-45-30-11 Eppendorf, Hamburg, Germany) the supernatant was passed over a C18 cartridge (500 mg, Varian, Middelburg, the Netherlands) to retain pigments. The effluent was then diluted to 50% methanol and concentrated on a DEAE-Sephadex (2ml, GE Healthcare Bio-Sciences AB, Uppsala, Sweden) anion exchange column for the analysis of free IAA and ABA, which are retained on the DEAE. The DEAE cartridge was eluted with 10ml 6% formic acid and IAA and ABA were concentrated on a C18 cartridge, which was coupled underneath. This C18 cartridge was eluted with 2×0.5 ml diethylether. The ether was evaporated under vacuum and the sample was dissolved in acidified methanol for methylation with diazomethane. After methylation, the samples were dried under a nitrogen stream and samples were further dissolved in 50 μl 10% MeOH for analysis.

IAA and ABA concentrations were determined following the protocol by Prinsen *et al*., (1995). Leaf and root IAA and ABA were analyzed by UPLC-MS/MS (Acquity TQD, Waters, Manchester, UK). The settings for the analysis were as follows: 6 μl injection by partial loop, column T. 30°C, solvent gradient 0-2 min: 95/5; 10% MeOH in NH4Oac 1 mM/MeOH; 2-4 min linear gradient until 10/90 10% MeOH in NH4Oac 1 mM/MeOH; 4-6 min, isocratic 10/90 10% MeOH in NH4Oac 1mM/MeOH; MS conditions: Polarity MS ES(+), capillary 2 kV, cone 20 V, collision energy: 20 eV, source temperature: 120 °C, desolvation Temperature: 450 °C, Cone gas flow 50 l/h, desolvation gas flow: 750 l/h, collision gas flow: 0.19 ml/h). The diagnostic ions used for quantification are 190>130 m/z for Me-IAA, 196>136 m/z for Me-C13-IAA, 279>173 m/z for Me-ABA and 285>179 m/z for d6-Me-ABA (dwell time 0.020 sec). Methanol and water used for MS are UPLC grade from Biosolve (Valkenswaard, the Netherlands). Data are expressed in pmol per gram fresh weight (pmol g^−1^ FW). Using this method, root ABA concentrations were too low to be detected since there were below the detection threshold of 10 pmol g^−1^ FW (Prinsen *et al*., 1995).

### Leaf anatomical measurements

During experiment E4, 5-cm leaf segments from the 8 selected genotypes were examined for leaf anatomical traits. These segments were carefully sectioned from the middle part (approx. 10 cm from the leaf tip) of the top, most fully developed leaf using a scalpel blade and instantaneously fixed with FAA solution [Ethanol 99% (45%vol), demineralized water (45%vol), Formaldehyde 36% (5%vol), acetic acid (5%vol)] during one week. Afterwards, they were transferred into a bleaching solution [Ethanol 99% (70%vol), acetic acid (30%vol)] for 3 days. Samples were then stored for approx. 3 weeks in an ethanol/water solution (70%vol / 30%vol) prior to further examination.

Freehand sections were produced using sharp razor blade and stained with a safranin solution [0.5 g/L – Ethanol/Water (1/19)] for 10 seconds before being quickly rinsed with water followed by 100% ethanol to avoid an over-coloration. Afterwards, thin leaf slices were mounted on microscope blade at an amplification ×400. Approximately fifty pictures were taken for each leaf in order to cover the entire transversal section of each leaf. Using the Image-J plugin Mosaic-J, images were then sequentially placed, adjusted and merged in order to obtain one complete, large angle and high resolution image of the entire leaf transversal section [Supplementary Data Fig. S2].

Those images were then used to quantify the leaf traits reported on Supplementary Data Fig. S2. The following anatomical traits were then determined on each leaf section image using distance and area measurement tools from image-J software: Leaf width (LW, mm), major and minor vein densities (respectively VD_M_, VD_m_, mm^−1^), average distance between veins (DV, μm), average vein section area (VSA, μm^2^), average meta-xylem section area (MXA, μm^2^), average leaf thickness (LT, μm). LW was measured as the length of the straight line passing through all the vein centers from one border of the leaf to the other [Supplementary Data Fig. S2]. VD_M_, VD_m_ were calculated as the leaf width divided by the number of major and minor veins, respectively. DV was measured from the centre of the vein to the centre of the following one (irrespective of the type of the vein) and averaged on the whole leaf cross section. VSA is the area delimited by the mestome sheath. MXA was calculated as the averaged section area of the two main meta-xylem vessels in each major vein. LT was determined from the leaf thickness measured at all the veins and between all the veins [Supplementary Data Fig. S2]. Abaxial and adaxial stomata densities (mm^−2^, AB_SD and AD_SD, respectively) were previously determined for those genotypes at a similar leaf position in a previous study (Schoppach *et al*., 2016).

### Data and statistical analysis

All statistical analyses (regression analyses, correlation analyses, one-way ANOVAs) were carried out using GraphPad PRISM version 7.0b (GraphPad Software Inc., San Diego, CA, 2017).

## RESULTS

### Daytime root auxin levels correlate with daytime transpiration and plant hydraulic conductance measured independently

Daytime root IAA concentrations significantly correlated with all three hydraulic traits examined in the study, namely whole-plant daytime TR (Fig. 1A), K_Plant_ (Fig. 1B) and the previously characterized slopes of whole-plant TR responses to increasing VPD (Fig. 1C), determined in Schoppach *et al*. (2016). In all cases, the correlations were negative, with Pearson’s r values ranging from −0.75 (root IAA vs. slopes) to −0.9 (root IAA vs. daytime TR). The correlation analysis also revealed that K_Plant_ strongly and positively correlated with the independently measured slopes of TR responses to VPD (Pearsons’s r = 0.84, P = 0.008, R^2^ = 0.72, Fig. 2), indicating that the selected 8 genotypes consistently exhibit the same hydraulic properties independent of the experiment.

**Fig. 1.**
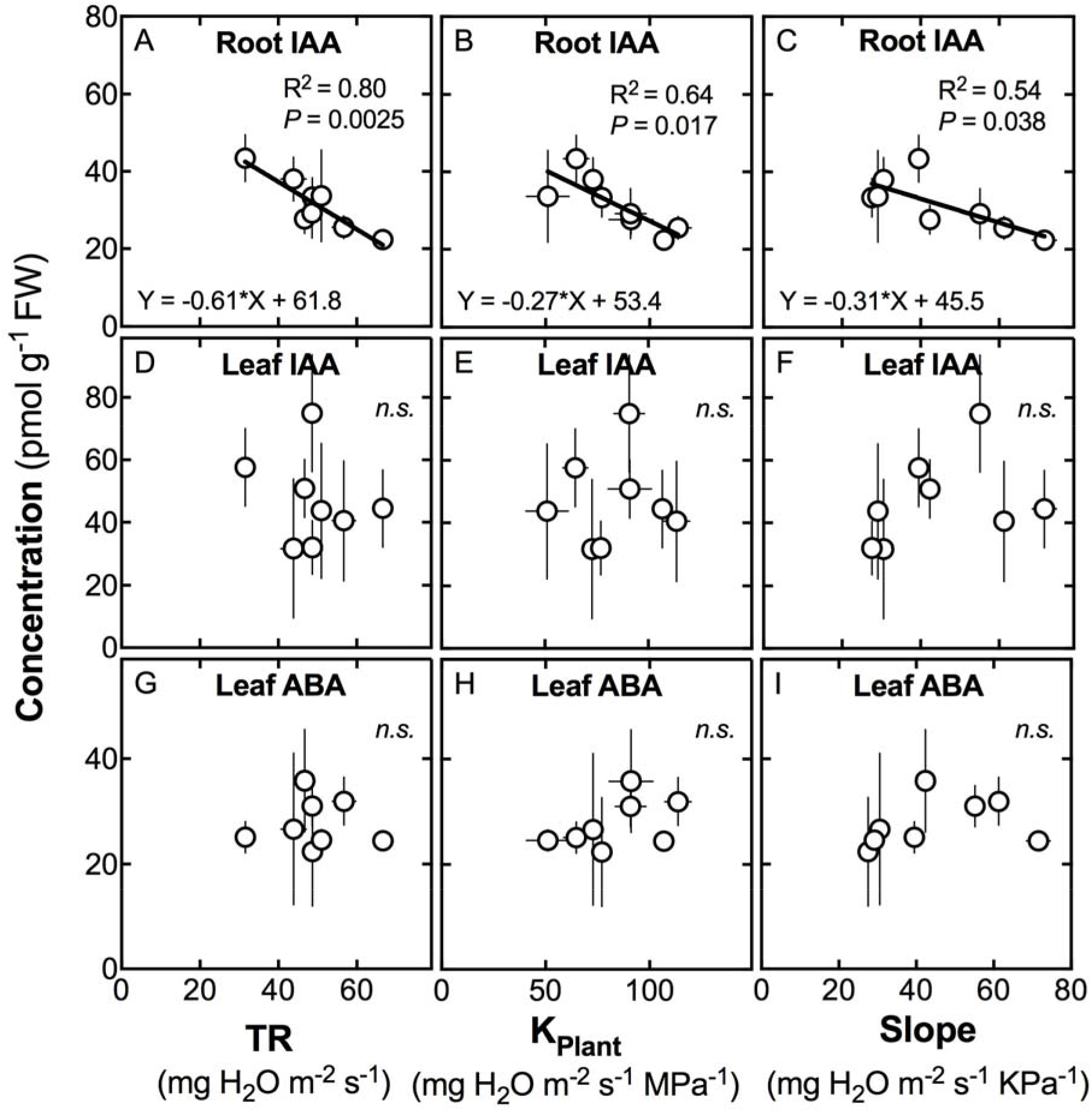
Relationships between root or shoot concentrations in indole acetic acid (IAA) or abscisic acid (ABA) and whole-plant daytime hydraulic traits in wheat. Panels A, B and C represent correlations between root IAA and whole plant transpiration rate (TR), plant hydraulic conductance (K_Plant_) and the slope of TR response to VPD, respectively. Panels D-F and G-I represent correlations between leaf IAA and leaf ABA concentrations and these same variables, respectively. When significant, statistical data and regression coefficients are indicated. n.s.: non-significant correlation. Each datapoint is the average of 3 observations (± S.E.) made on 8 double haploid wheat lines resulting from the cross between parents RAC875 and Kukri.

**Fig. 2.**
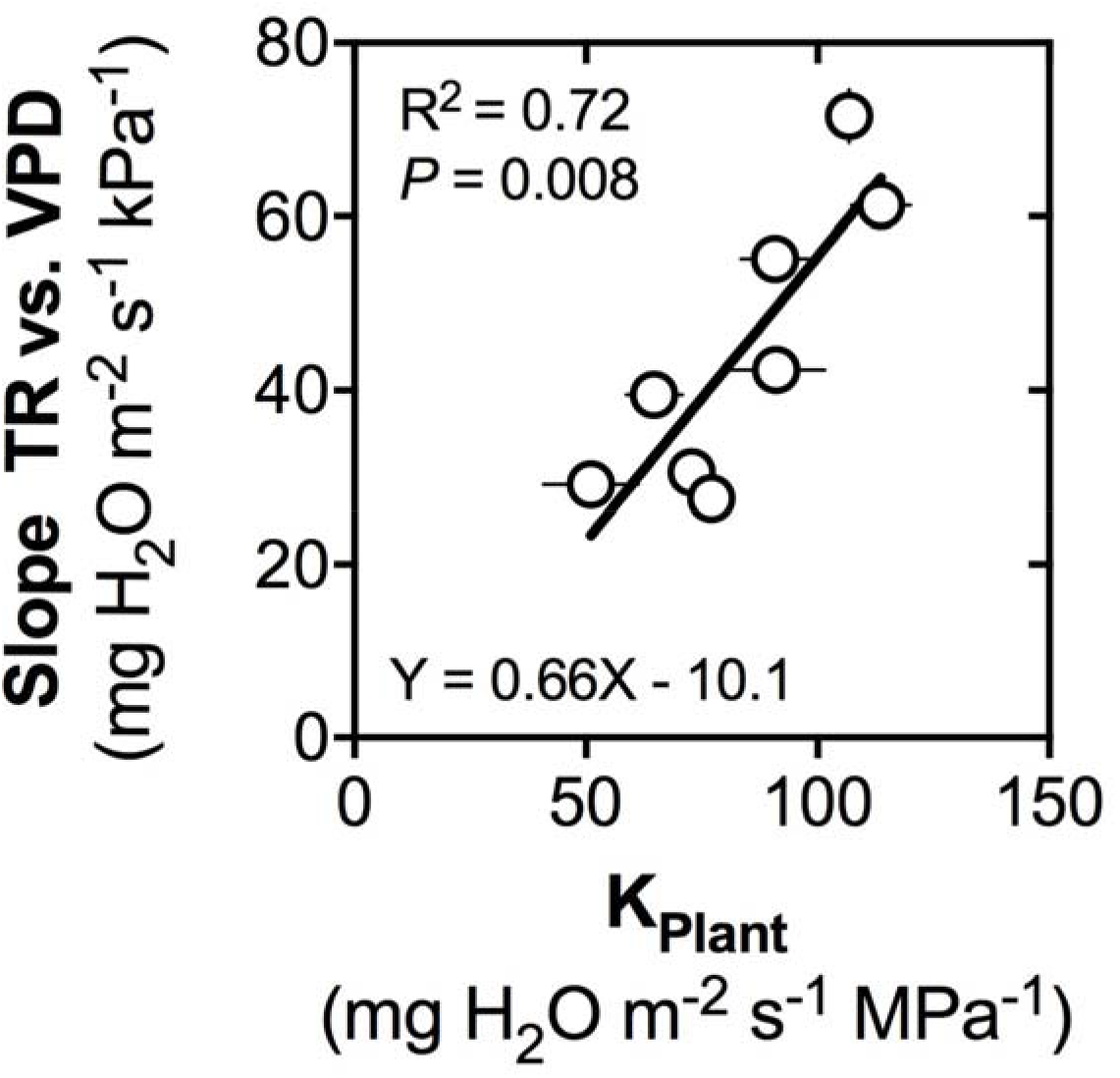
Relationship between the slope of whole-plant transpiration rate (TR) response curve to increasing vapor pressure deficit (VPD) determined in Schoppach *et al*., 2016) and plant hydraulic conductance (K_Plant_) measured independently in experiment E1 on the 8 wheat genotypes of the study.

In sharp contrast to daytime root IAA, daytime leaf IAA and leaf ABA concentrations were found to not significantly correlate with daytime whole-plant TR, K_Plant_, or the slopes of whole-plant TR responses to increasing VPD (Figs. 1D-1I). Further, daytime leaf IAA and leaf ABA concentrations did not correlate with any of the examined leaf vascular traits.

### Night-time root IAA levels are stable across experiments

Regardless of the time of the day, root ABA concentrations were below the detection levels of the method used for quantification (see Materials and Methods). Irrespective of the genotype, night-time leaf IAA, root IAA and leaf ABA concentrations did not vary significantly between experiments E3.1 and E3.2 (Fig. 3A). Experiment E2 exhibited significantly lower leaf IAA and higher leaf ABA concentrations (Fig. 3B), but in contrast, night-time IAA root concentrations were more stable and did not exhibit significant variation across all 3 experiments (Fig. 3C). Correlation analyses showed that genotypic rankings in night-time leaf IAA and leaf ABA levels were not consistent across experiments, even between E3.1 and E3.2, and none of these concentrations significantly correlated with leaf vasculature measurements.

**Fig. 3.**
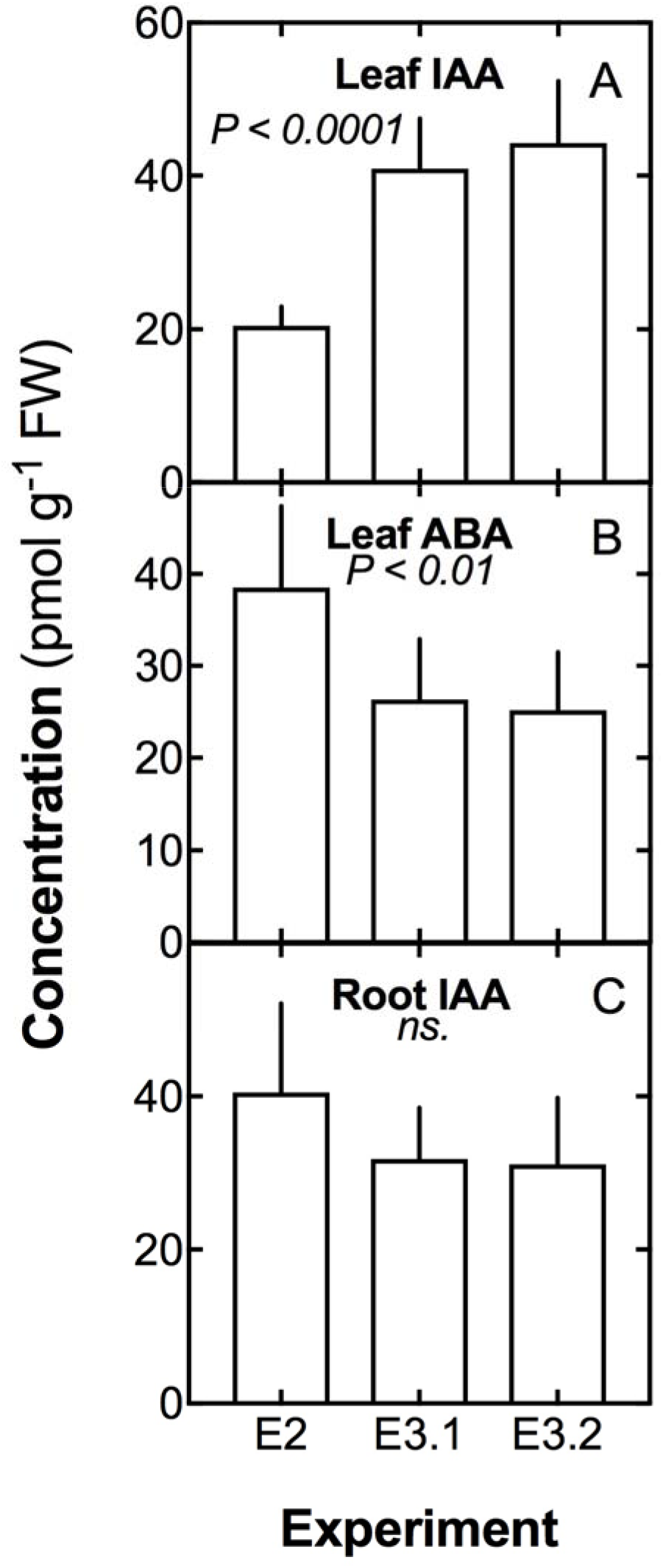
Variation in nighttime leaf concentrations in indole acetic acid (IAA, panel A), nighttime abscisic acid (ABA, panel B) and in nighttime IAA root concentrations (panel C) across 3 experiments (see Tables 1 and 2 for details about environmental conditions).

Root IAA concentrations were highly correlated between E2 and E3.1 (r = 0.97, P < 0.0001) and between E3.1 and E3.2 (Pearson’s r = 0.74, P = 0.037), while the correlation between E2 and E3.2 was significant at P = 0.08 (Pearson’s r = 0.65) as a result of an outlier datum (P < 0.05 if removed). Therefore, night-time root IAA data was pooled across the 3 experiments for further analysis (see below).

### Night-time root IAA is genotype-dependent and correlates strongly with daytime root IAA and with nocturnal transpiration rates

As reported on Fig. 4, variation in root IAA was found to be strongly dependent on genotypes both during the night (P<0.0001, Fig. 4A) and the day (P<0.005, Fig. 4B). During the night, root IAA concentrations ranged from 24.3 pmol g^−1^ for genotype DH_127 to more than twice that value, around 49.8 pmol g^−1^ for genotype DH_035. This variation was spanning a similar range during the day, from 22.4 pmol g^−1^ (genotype DH_127) to 43.4 pmol g^−1^ (genotype DH_003), with a similar ranking among genotypes (compare panels A and B in Fig. 4). Consistently, root IAA concentrations were found to be strongly and positively correlated among these 8 lines (Pearson’s r = 0.83, R^2^ = 0.68, P = 0.011, Fig. 5).

**Fig. 4.**
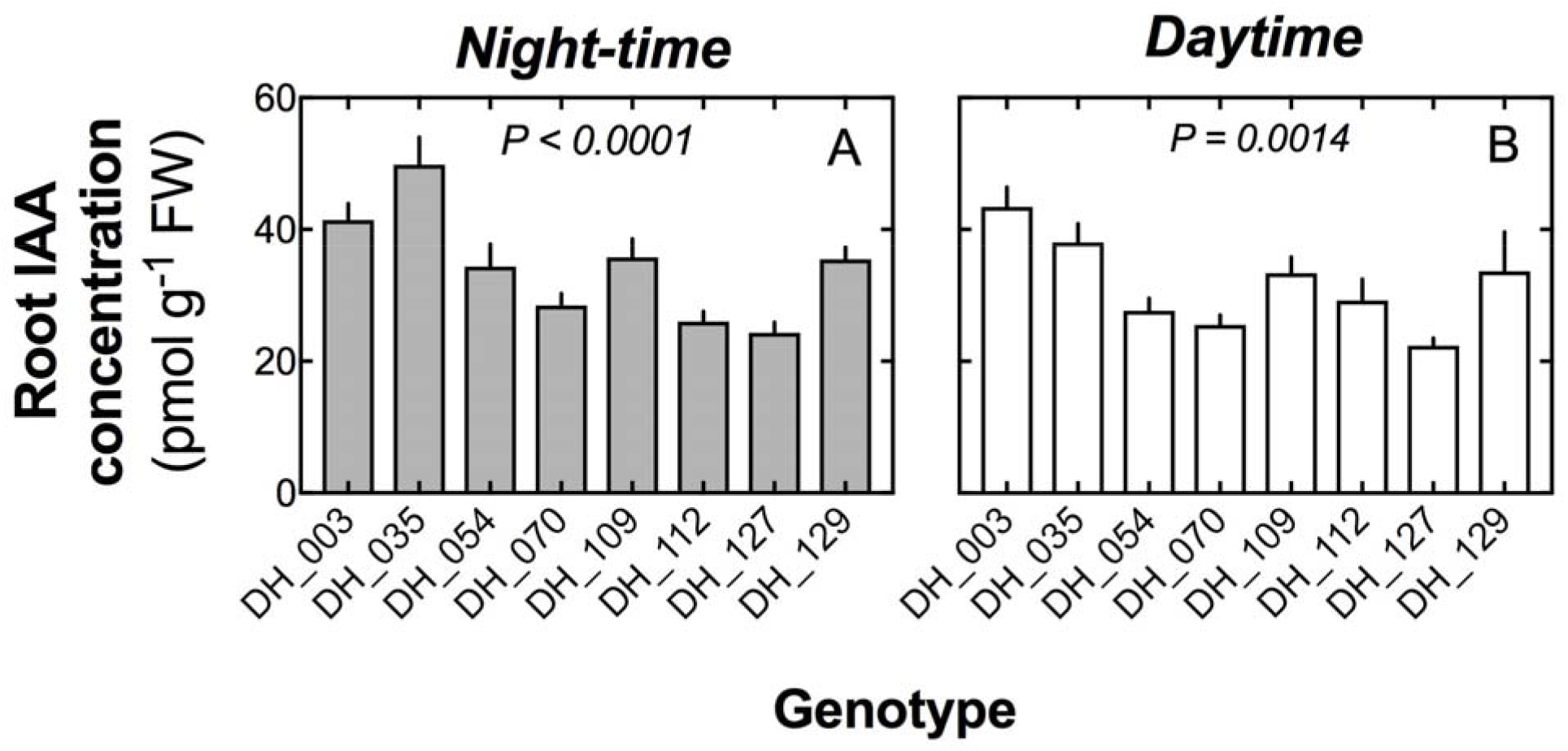
Genetic variability in night-time (grey bars) and daytime (open bars) indole acetic acid (IAA) concentrations in the roots of the 8 wheat genotypes of the study. P-values are significance levels of one-way ANOVAs testing for the genotypic effect of the differences in means between genotypes. Number of observations per genotype ranged from 4 (daytime) to 12 (night-time).

**Fig. 5.**
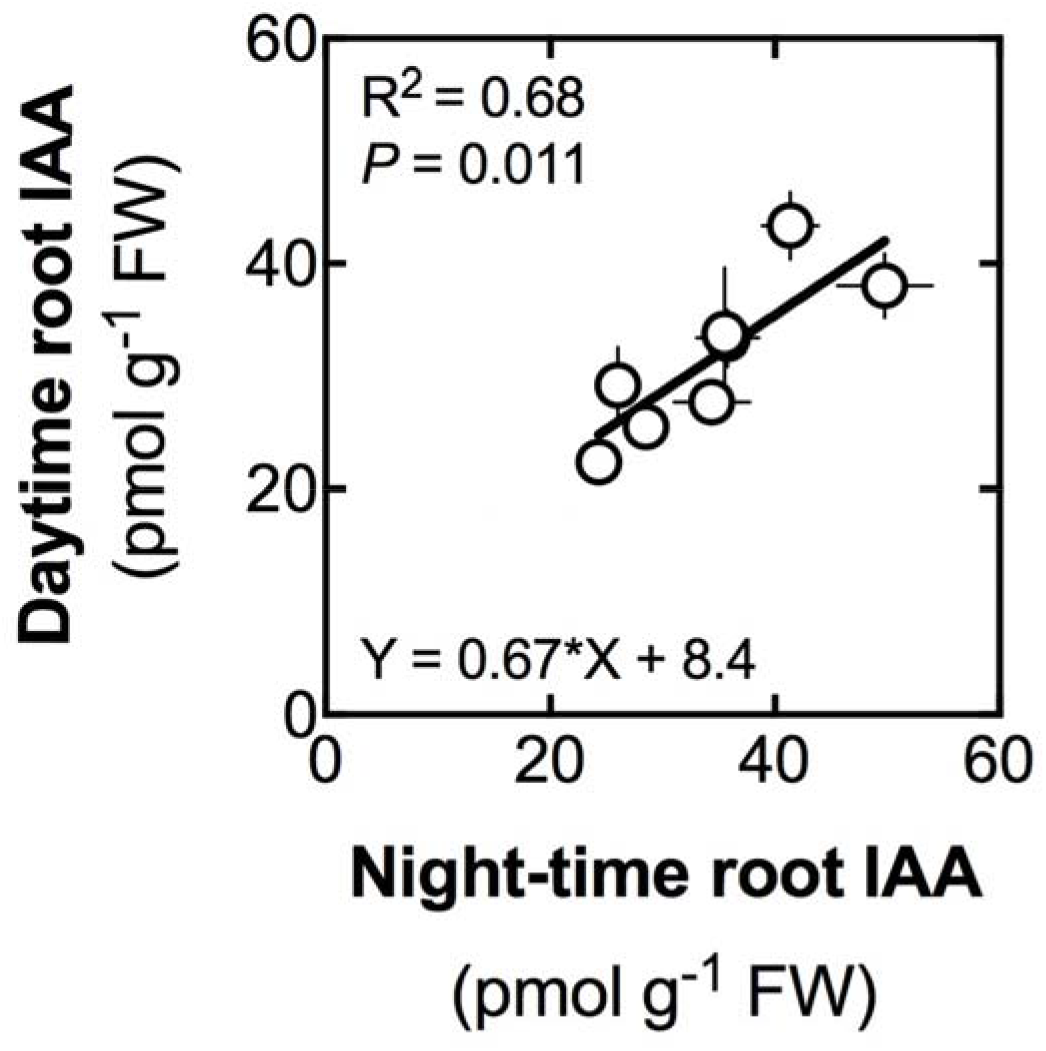
Relationship between nocturnal and daytime indole acetic acid (IAA) concentrations in the roots of the 8 studied wheat genotypes. Each datapoint represents a given genotype. Daytime and nighttime values are the average of 4 and 12 observations, respectively. Statistical data (R^2^, P-value and regression coefficients) are indicated.

Regardless of the experiment, night-time root IAA concentrations correlated significantly with TR_N_ (Pearson’s r = −0.69, R^2^ = 0.48, P < 0.0005, Fig. 6). This correlation was confirmed (i.e., statistically significant) when analysing separately data from experiments E3.1 and E3.2 (not shown), while experiment E2 displayed a very similar but non-significant tendency as a result of one single outlier datum (R^2^ = 0.81 and P < 0.01 if outlier not included). Night-time root IAA concentrations did not correlate with any of the leaf anatomical traits examined.

**Fig. 6.**
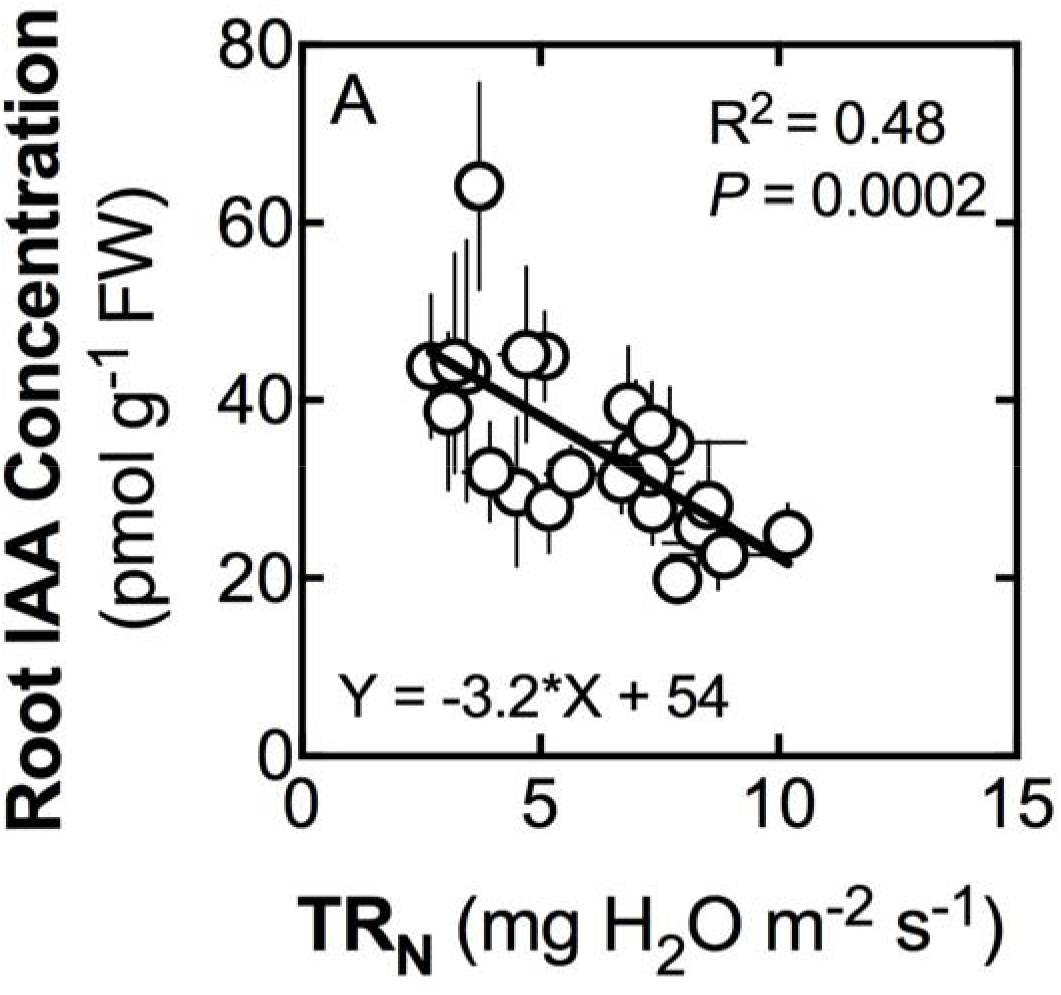
Relationship between whole-plant nocturnal transpiration rate (TR_N_) and root concentrations in indole acetic acid (IAA) in among the 8 studied wheat genotypes. Each datapoint is the average of 3-4 (± S.E.) observations. Data is pooled from three independent experiments (see Materials and Methods for details). Statistical data for the regression (R^2^, P-value and regression coefficient) are indicated.

## DISCUSSION

### Root auxins are potentially involved in regulating daytime whole-plant hydraulics

A first major finding of this research was that root IAA –but not leaf IAA– was potentially involved in controlling daytime water use in wheat, in a way that is consistent with the contrasted root hydraulic properties of the parents of the studied population and with what we already know about auxin putative roles in plant hydraulics (see below for details). To our knowledge, this is the first time that auxin is shown to be involved in whole-plant hydraulics, with a role in the expression of a water saving-trait that has been mechanistically linked to improved yields under terminal drought conditions.

In this investigation, this finding stems from the convergence of three independent sources of evidence, highlighting a negative correlation between root IAA and i) whole-plant TR (Fig. 1A), ii) K_Plant_ (Fig. 1B) and iii) the slope of TR response curves to increasing VPD (Fig. 1C). In all cases, those correlations were of the same sign, collectively indicating a putative involvement of auxin accumulation in the root in the expression of a hydraulic restriction at the canopy level. Such results are in line with Péret *et al*. (2012) who found that accumulation of root auxin decreases hydraulic conductivity at both the cell and whole root levels by negatively regulating AQP expression, in support of previous findings of Paciorek *et al*. (2005) who demonstrated that auxin suppress the endocytosis of PIP2 aquaporin in *Arabidopsis thaliana*.

These findings also strongly support previous results established on the parents of this population which indicated that lower TR under high VPD expressed by the drought-tolerant parent RAC875 relative to the drought-sensitive parent (Kukri) is likely stemming from a lower hydraulic conductivity of the roots of the former. Interestingly, this hydraulic limitation was found to be associated with decreased CMX vessel diameters combined with a restriction on the radial, AQP-mediated water transport in the root (Schoppach *et al*., 2014a)-- both of which are influenced by auxin. Indeed, local accumulation of root auxin is consistent with their role in repressing of root aquaporins (Péret *et al*., 2012) and in decreasing the vascular cell size (Fàbregas *et al*., 2015). Furthermore, the involvement of root auxin is consistent with the outcome of the QTL mapping carried out on the DH population from which the 8 genotypes of the study were selected, and the independent RNASeq analysis which mapped the peak region of the major QTL for TR responses to VPD to root-specific transcripts with functions suggesting involvement of root auxin (Schoppach *et al*., 2016).

The lack of correlation between leaf IAA, leaf ABA and the hydraulic variables examined in this study seems to further support the idea of a predominant involvement of auxin-mediated root hydraulic processes in controlling whole-plant water use in wheat, at least in this population. However, our inability to measure root ABA in this study does not allow to discard the possibility that genotypic variation in root ABA would have contributed to variation in TR response to VPD, as suggested by a previous study on durum wheat (Kudoyarova *et al*., 2011). Furthermore, our study does not discard the possibility that auxin effects were mediated through interaction with other plant hormones such as cytokinin and ethylene as suggested by studies highlighting such cross-talks (Tanaka *et al*., 2006; Rowe *et al*., 2016). Regardless, the above findings make it clear that root auxin plays an important role in regulating whole-plant water use and hydraulic properties at least in this population.

### A potential role for root auxins in controlling nocturnal transpiration

The second major finding of this investigation was that nocturnal water use is strongly and negatively correlated with auxin accumulation levels in wheat roots (Fig. 6). This finding was consistently observed over three independent experiments. To our knowledge, this is the first time that variation in hormonal levels were associated with changes in nocturnal water use. Importantly, the negative correlation between root auxin concentrations and TR_N_ was consistent with the relationship linking daytime root auxin concentration and daytime water use, indicating that root auxin accumulation tends to reduce transpiration regardless of the time of the day, a hypothesis reinforced by the strong correlation between daytime and night-time auxin levels in the roots (Fig. 5). This in turn may help explain why the drought-tolerant parent RAC875 expressed the water-saving behaviour both during the day and the night, and more generally, the previously reported strong correlation between daytime and night-time slopes of TR response to VPD on wheat (Schoppach *et al*., 2014b). Similar to daytime conditions, leaf IAA, leaf ABA and leaf vascular traits were not found to correlate with TR_N_ values, indicating that this TR_N_ variation in this population is largely controlled by roots. However, as previously stated, the additional involvement of root ABA could not be discarded, given that the root ABA concentrations were below the detection levels of the method used for quantification.

The dominating theories explaining the functional relevance of nocturnal water use typically revolve around transport mechanisms, defining a trade-off space where roots are central players. For instance, it is thought that increased TR_N_ would facilitate nitrogen acquisition by roots grown in N-limited environments (Rohula *et al*., 2014), and oxygen delivery in the xylem sap to sustain dark respiration-mediated nocturnal carbohydrate export (Marks and Lechowicz, 2007). On the other hand, increases in TR_N_ could also decrease the rate of hydraulic redistribution in the soil, a phenomenon driving the movement of water from moist to dry soil through the root system, which was associated with enhanced daytime transpiration efficiency and whole-season growth (Howard *et al*., 2009; Neumann *et al*., 2014). In concert with studies documenting regulatory effects of root auxin on aquaporin-mediated root hydraulic conductivity (Péret *et al*., 2012), xylem vessel development (Fàbregas *et al*., 2015), and nitrogen foraging by the roots (Song *et al*., 2013), our results indicate that the root auxins are potentially involved in the trade-offs associated with TR_N_.

### Root auxins: implications for drought tolerance

The key findings of this research are consistent with a stream of recent publications suggesting the involvement of root auxins in crop drought tolerance. Our own findings on wheat indicate that root auxin accumulation potentially drives the restriction in root hydraulic conductivity that is associated with the expression of the water-saving limitation of whole-plant TR, particularly under times of the day with high evaporative demand (Schoppach *et al*., 2014a) or the night (Schoppach *et al*., 2014b, Claverie *et al*., 2018). Other research suggests yield benefits resulting from auxin accumulation in the roots, although the link with whole-plant hydraulics was not documented. On maize, Li *et al*. (2018) found that maize mutants over-expressing ZmPIN1a exhibited an increase in auxin export from the shoot to the root where its accumulation in the root tips drove enhanced root development that was associated with increased yields under drought, likely through improved water acquisition. On wheat, the over-expression of the gene Auxin Biosynthetic TRYPTOPHAN AMINOTRANSFERASE RELATED TaTAR2.1-3A was associated with enhanced grain yield under various N supply levels that was likely to be the result of enhanced lateral root branching that promoted N foraging capabilities. Importantly, this gene was found to be predominantly expressed in the roots and its overexpression generated elevated auxin accumulation in the primary and lateral root tips (Shao *et al*., 2017). Finally, on sorghum, multiple QTL controlling stay-green, a trait that is linked to tolerance to terminal drought, have been found to co-localize with markers associated with an auxin-responsive gene (indole-3-acetic acid-amido synthetase GH3.5, Rama Reddy *et al*., 2014) or to genes from the PIN family of auxin efflux carriers (Borrell *et al*. 2015).

Taken together, these results indicate root auxin accumulation drive yield increases under drought either via a dynamic regulation of root hydraulic conductivity in order to express a water-saving behavior or via a developmental control of branching to optimize water capture, two strategies that have been proven to be widely effective in drought breeding (e.g., Vadez 2014; Wasson *et al*., 2012).

### Caveats and limits

This study suggests an important role for root IAA in controlling daytime and nocturnal water use, but does not necessarily discard the hypothesis that root and shoot ABA are also involved in regulating whole-plant TR under increasing VPD as previously found on durum wheat (Kudoyarova *et al*., 2011). Indeed, in this investigation, root ABA levels were below the detection threshold of the method used for quantification, which is around 10 pmol g^−1^ FW, while in the above study, nominal root ABA concentrations were equivalent to 6.8 pmol g^−1^ FW. However, it is noteworthy that in Kudoyarova *et al*. (2011), experiments were carried on another species, (durum wheat), on young seedlings (7-day old) and using a different technique (immunoassay). While always possible, an error in the protocol used in this study is unlikely, considering the consistency of lack of detection of root ABA and the consistent levels of root IAA across independent experiments and genotypes.

Another potentially confounding effect in this research was that leaf-level determinations of IAA and ABA concentrations were made on the basis of single leaves, while root concentrations were made on the basis of the bulk root system. While we harvested tissue segments from leaves having similar age and positions in upper layer of the canopy, it is possible that leaf-to-leaf variation masked correlations between leaf auxin or ABA and the examined traits. That said, whole-root system hormonal dosage does not necessarily guarantee stable, consistent measurements, since roots of different ranks, age and microenvironment are not necessarily expected to exhibit the same hormonal concentrations. In any case, further studies are required to illuminate i) the exact mechanisms controlling auxin redistribution from the shoot to the root, ii) the interplay between root auxin accumulation and root/shoot hydraulic properties and iii) the role played by interactions with other hormones such as ABA, ethylene and cytokinin.

## CONCLUSIONS

This study indicates that variation in whole-plant transpiration responses to evaporative demand, a major trait that drives the expression of water-saving, drought-tolerance strategies in wheat, is potentially controlled by auxin levels in the root system. Specifically, root auxin accumulation was found to be negatively correlated with instantaneous TR, the slope of whole plant TR response to VPD and plant hydraulic conductance. Furthermore, we unravelled a previously undocumented association between root auxin and nocturnal water use in a way that is consistent with its role as a negative regulator of hydraulic conductance.

Those findings illuminate potentially important roles of root auxin in regulating daytime and nocturnal water use that are consistent with previous physiological, genetic and molecular evidence established on the studied population. They are also in line with evidence from other sources documenting a role of root auxins in regulating hydraulic conductivity and enhancing crop yields under water-limited environments. While this study suggests that root auxin levels might be a stable trait that could predict drought tolerance capabilities of wheat genotypes, further investigation is needed to mechanistically link auxin accumulation and its hydraulic and developmental consequences at the local and the whole-plant levels and in relation to other hormones.

## ACKNOWLEDGEMENTS

We thank Elisabeth Majerus for her help in preparing leaf sections and Coline Kinkin for her help in grinding tissue samples for hormone dosage.

---

**Supplementary data fig. S1.**
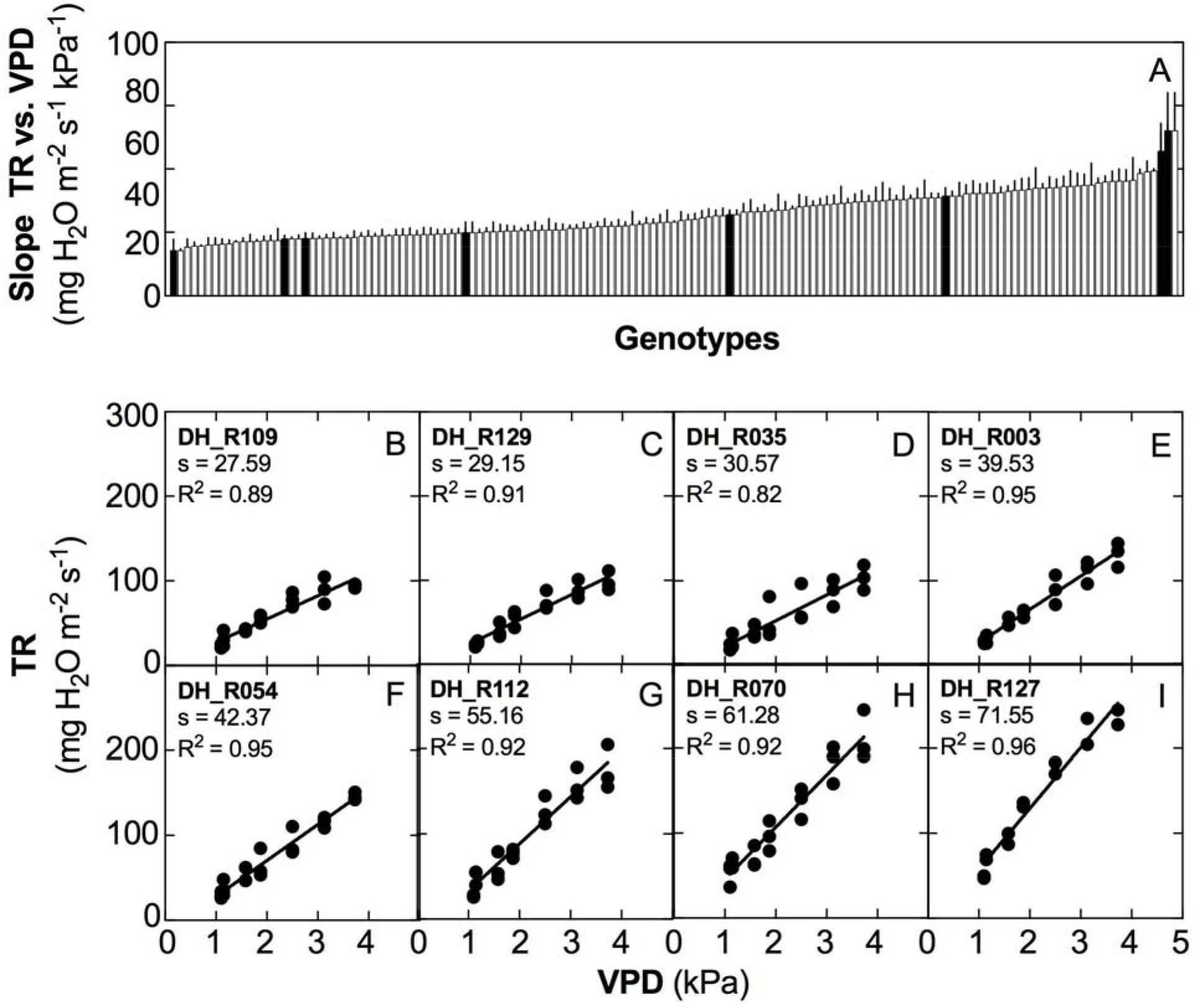
Diversity in the slopes of whole-plant transpiration rate (TR) response curves to naturally increasing vapor pressure deficit (VPD) within the population of 143 double haploid (DH) lines from the RAC875 x Kukri cross (panel A) and among the 8 lines selected lines (panels B-I). Panel A: bars representing the values of the slopes (± S.E.) are ordered from the lowest to the highest. Black bars in panel A are for the 8 lines that were selected. Panels B to I represent the TR response curves to increasing VPD for each of the lines where the slopes of the linear regressions (s) and the coefficients of determination (R^2^) are reported. Data is from Schoppach *et al*., (2016).

**Supplementary data fig. S2.**
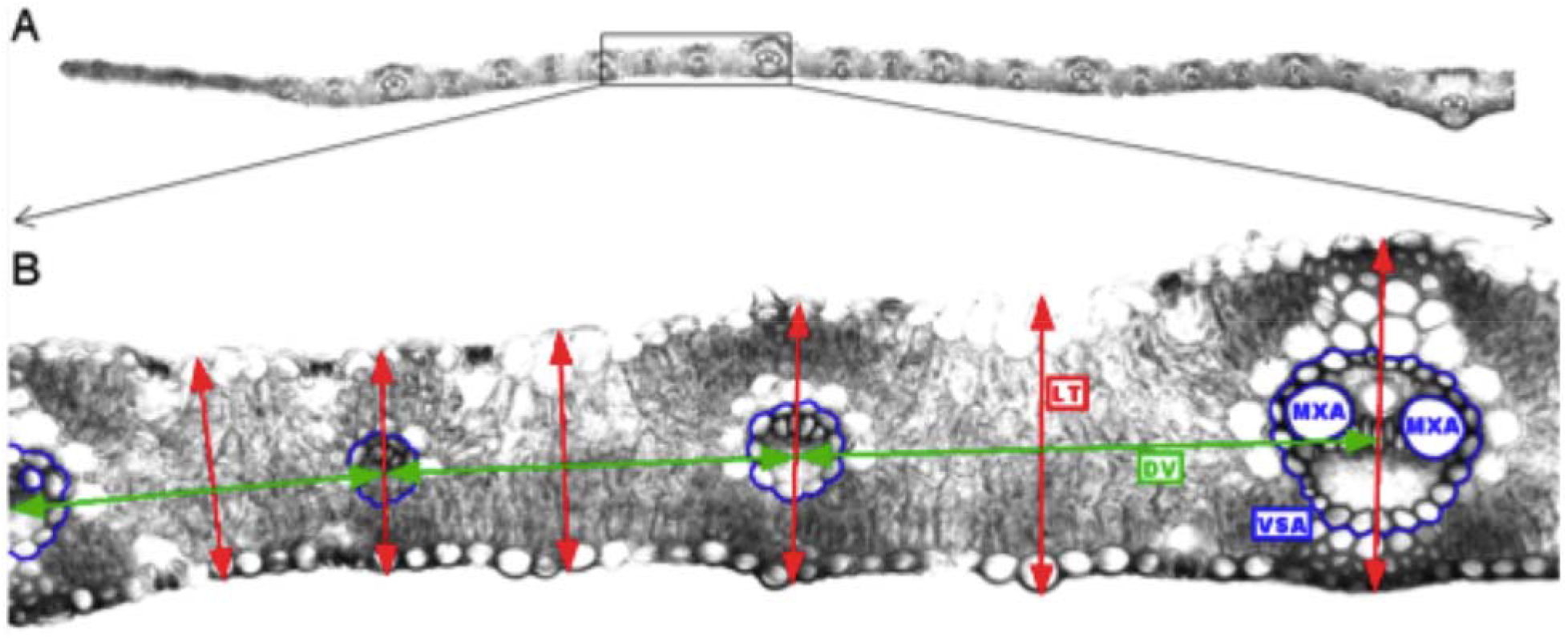
Illustration of the examined leaf anatomical traits among the 8 lines of the study. Panel (A) represents a composite image based on the merger of 48 pictures to reconstruct a transversal view of the entire leaf. Only 50% of the image is presented here for convenience. Panel (B) represent a zoom of an area from the top picture and illustrates the traits of interest of this investigation. Blue lines delineate the vein section area (VSA) and the two main meta-xylem vessels areas (MXA) in each major vein. Horizontal arrows represent the distances (from the center) between two successive veins, which were used to calculate leaf width (LW), vein densities (VD_M_ and VD_m_) and average distance between veins (DV). Vertical arrows represent the leaf thickness measured at each position (at each vein and midway between consecutive veins) and averaged over the entire leaf to calculate average leaf thickness (LT, see Material and Methods for details).

